# DNA methylation directs polycomb-dependent 3D genome reorganisation in naïve pluripotency

**DOI:** 10.1101/527309

**Authors:** Katy A. McLaughlin, Ilya M. Flyamer, John P. Thomson, Heidi K. Mjoseng, Ruchi Shukla, Iain Williamson, Graeme R. Grimes, Robert S. Illingworth, Ian R. Adams, Sari Pennings, Richard R. Meehan, Wendy A. Bickmore

**Affiliations:** MRC Human Genetics Unit, Institute of Genetics and Molecular Medicine, University of Edinburgh, Crewe Road South, Edinburgh EH4 2XU, UK; Northern Institute for Cancer Research, Framlington Place, Medical Faculty Newcastle upon Tyne, NE2 4HH, UK; Centre for Cardiovascular Science, Queen’s Medical Research Institute, University of Edinburgh, 47 Little France Crescent, EH16 4TJ, UK

**Keywords:** 3D genome, DNA methylation, fluorescence in situ hybridisation, Hi-C, pluripotency, polycomb

## Abstract

The DNA hypomethylation that occurs when embryonic stem cells (ESCs) are directed to the ground state of naïve pluripotency by culturing in 2i conditions results in redistribution of polycomb (H3K27me3) away from its target loci. Here we demonstrate that 3D genome organisation is also altered in 2i. We found chromatin decompaction at polycomb target loci as well as loss of long-range polycomb interactions. By preventing DNA hypomethylation during the transition to the ground-state, we are able to restore the H3K27me3 distribution, and polycomb-mediated 3D genome organisation that is characteristic of primed ESCs grown in serum, to ESCs in 2i. However, these cells retain the functional characteristics of 2i ground state ESCs. Our findings demonstrate the central role of DNA methylation in shaping major aspects of 3D genome organisation but caution against assuming causal roles for the epigenome and 3D genome in gene regulation and function in ESCs.

## Introduction

The extent to which epigenetic modifications and three-dimensional chromatin structure are linked and contribute to cell state and cell function is unresolved. Two key and inter-related epigenetic modifications in the mammalian genome are DNA methylation and histone modification by polycomb group proteins. Polycomb complexes are implicated in the maintenance of repression of key developmental genes (Blackledge et al., 2015). Whereas polycomb repressive complex PRC2 deposits H3K27me3, the canonical PRC1 complex promotes compact local chromatin structures and longer range chromatin interactions (Boettiger et al., 2016; Eskeland et al., 2010; Joshi et al., 2015; Kundu et al., 2017; Schoenfelder et al., 2015; Williamson et al., 2012). Chromatin compaction and developmental gene repression are independent of the E3 ligase catalytic activity of Ring1B in canonical PRC1 (Cohen et al., 2018; Eskeland et al., 2010; Illingworth et al., 2015; Kundu et al., 2017; Williamson et al., 2014).

In mammalian cells, the polycomb system is primarily targeted to the unmethylated CpG islands (CGI) of non- or weakly-expressed genes (Blackledge et al., 2015; Li et al., 2017; Perino et al., 2018; Riising et al., 2014). Consistent with this, loss of DNA methylation - by exposing new CpG sites - leads to a redistribution of H3K27me3 across the genome, to satellite and dispersed repeat sequences, and titrating it away from its normal CGI targets (Brinkman et al., 2012; Jermann et al., 2014; Reddington et al., 2013; Reddington et al., 2014). This is consistent with a model in which PRC2 can associate transiently and weakly with a large fraction of the genome (Schuettengruber et al., 2017).

One notable instance in which this occurs is in mouse embryonic stem cells (mESCs) cultured under 2i conditions (Marks et al., 2012). mESCs cultured conventionally in the presence of foetal calf serum and LIF are functionally heterogeneous, with a fraction of cells resembling a state of ‘naive pluripotency’ - with unbiased developmental potential and high expression of pluripotency genes. Other cells more closely resemble a “primed” state in which cells begin expressing early lineage markers and down regulate pluripotency genes (Canham et al., 2010; Hackett and Surani, 2014; Hayashi et al., 2008; Wongtawan et al., 2011). These are two metastable states between which the cells in the population fluctuate. By contrast, culturing mESCs serum-free, in the presence of two small molecule inhibitors (2i) of MEK1 (also called MKK1 (MAPK kinase-1)) and glycogen synthase kinase 3 (GSK3), blocks differentiation signals and directly promotes the pluripotency network. This promotes naive pluripotency homogeneous expression of key pluripotency factors and reduced expression of early lineage-specific genes (Morgani et al., 2013; Silva and Smith, 2008; Wray et al., 2011; Ying and Smith, 2017).

Epigenetic properties of 2i cultured mESCs closely resemble those of cells in the pre-implantation inner cell mass (ICM) of the mouse embryo. This includes global DNA hypomethylation (Ficz et al., 2013; Leitch et al., 2013; Marks et al., 2012; Wray et al., 2011). Expression levels of the de novo methyltransferases Dnmt3a, 3b and the non-catalytic cofactor Dnmt3l are reduced under 2i conditions. Uhrf1 (a Dnmt1 co-factor) is also down-regulated at the protein level (Ficz et al., 2013; Grabole et al., 2013; Graf et al., 2017; Habibi et al., 2013; Leitch et al., 2013; von Meyenn et al., 2016; Yamaji et al., 2013). However, coupling these DNA methylation differences to gene expression changes using triple knock out (TKO) cells that lack all the active Dnmts, reveals that only a small (but significant) proportion of gene expression changes under 2i can be directly attributed to DNA methylation loss (Leitch et al., 2013).

Importantly, although global levels of H3K27me3 are not altered in 2i-cultured cells, there is a marked reduction (up to 75%) of H3K27me3 at polycomb targets - including at the Hox clusters (Marks et al., 2012). The consequences of such a dramatically altered epigenome on three-dimensional (3D) genome organisation have not been explored. Given the dynamics of epigenetic alterations that occur under 2i culturing conditions, and the role of polycomb in shaping the 3D genome, we sought to investigate whether 2i impacts on 3D chromatin organisation in mESCs. Using fluorescence in situ hybridisation (FISH) and Hi-C, we show that local chromatin compaction at polycomb-target Hox loci and long-range polycomb interactions are profoundly altered under 2i/LIF, and we demonstrate that this is directly attributable to the loss of DNA methylation. By restoring the epigenetic landscape (DNA methylation and H3K27me3 targeting) of cells in 2i/LIF we show that 3D genome organisation can be reset to resemble that of mESCs grown in serum/LIF. Strikingly, this has a limited impact on gene expression, despite having an epigenetic landscape and 3D organisation comparable to mESCs growing in serum/LIF.

## Results

### Chromatin decompaction of polycomb target loci in naïve ES cells

Mouse ESCs cultured in a chemically defined medium in the presence of LIF and two inhibitors (2i) of the Erk and Gsk-3 signalling pathways achieve a homogeneous ground state of pluripotency, thought to closely resemble that of the inner cell mass (ICM) (Ying et al., 2008; Ying and Smith, 2017). In doing so, 2i-mESCs acquire a distinct epigenomic landscape, including global DNA hypomethylation and an altered genomic distribution of H3K27me3 (Ficz et al., 2013; Habibi et al., 2013; Leitch et al., 2013; Marks et al., 2012) This includes a loss of H3K27me3 enrichment at classic polycomb targets such as Hox loci (Figure 1A).

**Figure 1:**
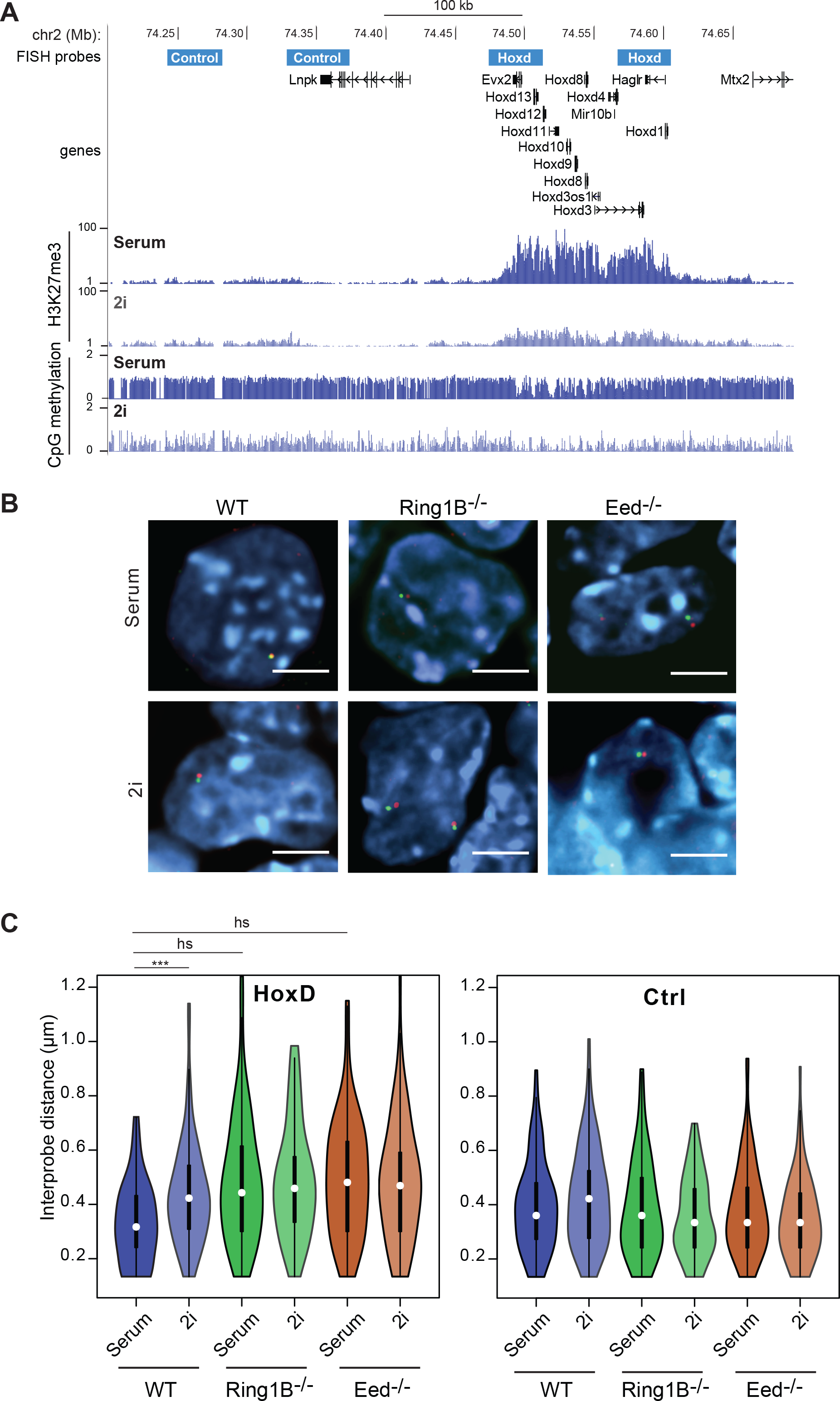
Loss of chromatin compaction at polycomb target loci in 2i. A. UCSC genome browser tracks (mm9 assembly) showing the location (Mb) on chromosome 2 of FISH probes used to measure compaction across the *HoxD* locus, and at a control locus. Probe co-ordinates are given in Supplementary Table 1. Below are shown the H3K27me3 ChiP-seq (Marks et al., 2012) and DNA methylation bisulphite (Habibi et al., 2013) profiles for this region of the genome in mESCs grown in serum or 2i. B. Representative images of FISH signals (red and green) from probes (indicated in A) detecting the *HoxD* locus in the nuclei of WT, *Ring1B*^−/−^ and *Eed*^−/−^ mESCs. DNA is counterstained with DAPI (blue). Scale bars represent 10 μm. C. Violin plots showing the distribution of inter-probe distances (μm) for *HoxD* and control (Ctrl) loci in WT, *Ring1B*^−/−^ and *Eed*^−/−^ cells grown in serum or 2i. The vertical line and spot within each plot indicate the interquartile range and median, respectively. ***p<0.001, h.s. highly significant (p<0.0001). Full details of statistical analysis are in Tables S2 and S3.

Since polycomb is a powerful mediator of higher-order chromatin structure (Boettiger et al., 2016; Eskeland et al., 2010; Francis et al., 2004; Joshi et al., 2015; Kundu et al., 2017; Schoenfelder et al., 2015; Williamson et al., 2012), it is possible that the redistribution of H3K27me3/polycomb across the genome results in an alteration to 3D chromatin organisation in ESCs grown in 2i culture conditions, but this has not been investigated.

The murine *HoxD* locus is a large canonical polycomb target in mESCs, demarked by a domain of H3K27me3, PRC2 and PRC1 deposition across the 100kb cluster (Illingworth et al., 2012). Under serum/LIF culture conditions the *HoxD* locus is maintained in a compact chromatin conformation in mESCs, and this is dependent on the presence of the PRC1 (Eskeland et al., 2010; Williamson et al., 2014). To investigate the degree of higher-order chromatin compaction at *HoxD* in mESCs grown under serum and 2i conditions, we used 3D FISH to measure the separation of hybridisation signals from probe pairs at opposite ends of the *HoxD* locus (*Hoxd3* and *Hoxd13*) under the different conditions. We compared these measurements to those from control probes in a nearby adjacent genomic region (3’ of *Lnp*), that is not coated by H3K27me3 but is highly DNA methylated in serum grown ESCs (Figure 1A).

FISH showed that, under 2i/LIF culture conditions, the *HoxD* locus significantly decompacts relative to cells cultured in serum/LIF; median inter-probe distances increase from ~300nm to ~400nm, p=<0.0001 (Figure 1B, C, Supplementary Figure 1A, Supplementary Tables 2 and 3). This decompaction occurs to the same extent when either PRC1 (*Ring1B* ^−/−^) or PRC2 (*Eed* ^−/−^) are absent in mESCs grown under serum conditions (Figure 1B, C, Supplementary Figure 1A). No further decompaction is observed when PRC1 or PRC2 mutant mESCs are grown under 2i conditions, showing that decompaction of a polycomb target in 2i can be primarily accounted for by the titration of H3K37me3/polycomb away from these genomic regions in 2i cells. We confirmed these data for two other Hox clusters – *HoxB* (Supplementary Figure 1B, D and F) and *HoxC* (Supplementary Figure 1C, E, G).

As a control, we examined a locus not marked by H3K27me3, and highly DNA methylated, in serum grown ES cells, that is adjacent to *HoxD* (Figure 1A). Inter-probe distances at this control locus were not significantly different between wild-type or polycomb mutant mESCs, or between mESCs grown in the different culture conditions (Figure 1C, Supplementary Figure 1A), even though this region is subject to DNA hypomethylation in 2i (Figure 1A). This suggests that the chromatin decompaction we detect in 2i conditions at polycomb target loci is not a result of a general/global alteration in the 3D chromatin organisation of naïve 2i/LIF cells, and that global loss of DNA methylation across genomic regions has no direct effect on chromatin compaction, as assayed at a cytological level.

### *HoxD* chromatin compaction in the blastocyst is comparable to that in 2i mESCs

Next, we investigated whether the chromatin decompaction observed in 2i-cultured mESCs is also present in the cells of the mouse blastocyst, which are naturally hypomethylated during normal development. To compare chromatin states between our *in vitro* mESCs and their *in vivo* counterparts, we measured distances between HoxD probes in E3.5 mouse blastocysts using 3D FISH (Figure 2A).

**Figure 2:**
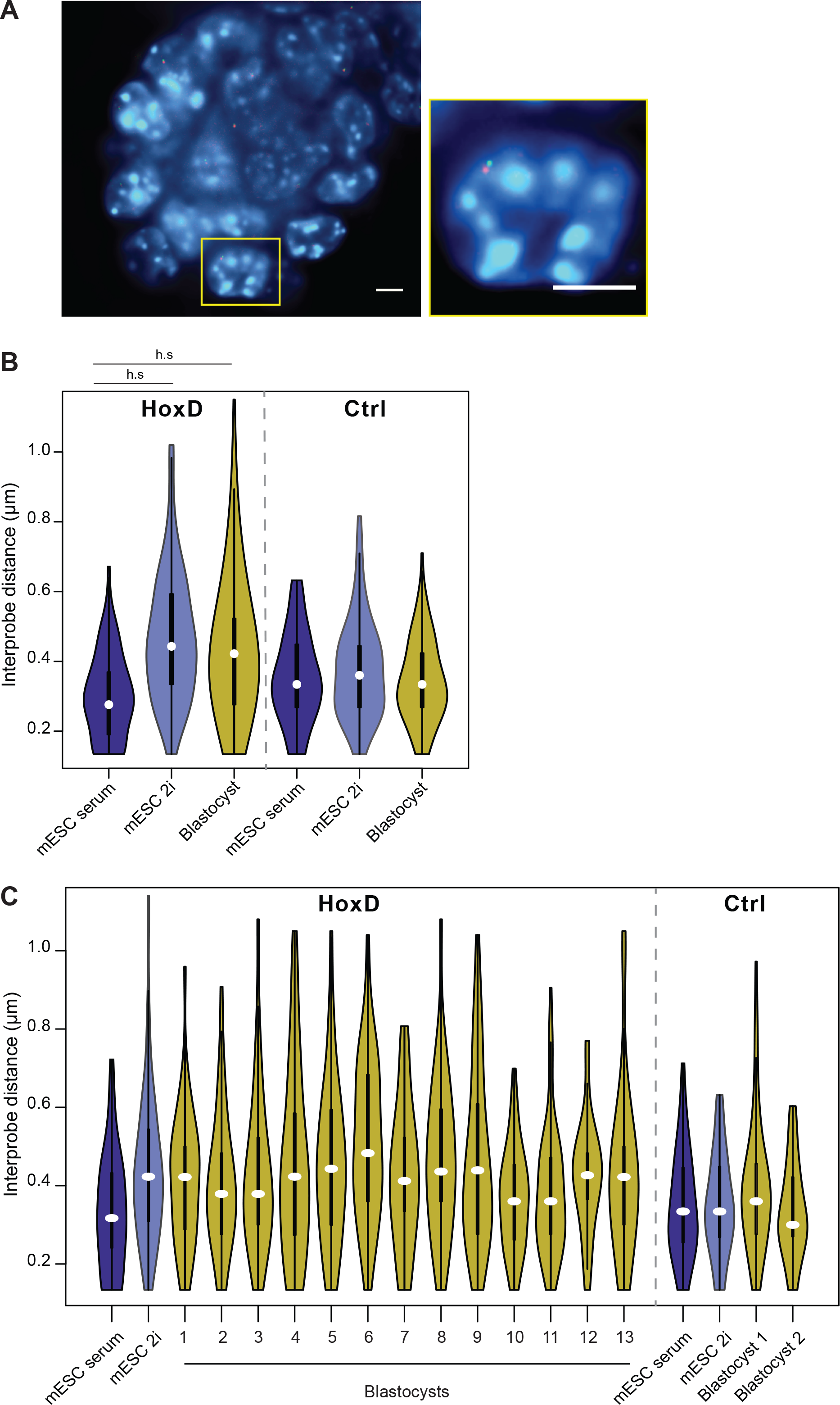
HoxD chromatin compaction in the pre-implantation blastocyst. A. Representative image of a DAPI-stained (blue) whole E3.5 blastocyst following FISH with probe pairs (red and green) detecting the *HoxD* locus. Inset shows enlargement of one nucleus. Scale bars represent 10 μm. B. Violin plots showing the distribution of inter-probe distances (μm) for *HoxD* and control (Ctrl) loci in E14 mESCs grown in serum or 2i, and in E3.5 blastocysts. Data are presented as in Figure 1C. h.s = p< 0.0001 C. As in (B) but data from 13 individual blastocysts.

These data indicate that the *HoxD* locus in the pre-implantation blastocyst is decompact relative to that in conventionally cultured serum/LIF mESCs, and closely resembles the compaction state of the locus under 2i/LIF conditions (Figure 2B and C, Supplementary Tables 2 and 3. There is a large amount of variability between and within blastocysts, which is likely because these blastocysts will contain distinct cell lineages (trophectoderm, ICM, and primitive endoderm), all of which are hypomethylated (Rossant et al., 1986). In contrast, inter-probe distances at the control locus were much more similar between blastocysts and cultured cells (Figure 2B), suggesting the decompaction at *HoxD* in the blastocyst cannot be explained by the *in vivo* population having a generally more open chromatin structure.

### Globally altered local interactions at polycomb loci between serum and 2i-cultured mESCs

Polycomb is responsible for forming self-interacting topologically associated domains (TADs) at *Hox* loci as detected by chromosome confirmation capture methods (Kundu et al., 2017; Noordermeer et al., 2011; Williamson et al., 2014). To assess whether changes in 3D chromatin organisation occur in 2i cells at regions other than *Hox* loci, we employed *in situ* Hi-C (Lieberman-Aiden et al., 2009; Rao et al., 2014) to assay genome-wide chromatin interactions from E14 mESCs grown in serum/LIF and in 2i/LIF. Visual inspection of the resulting contact frequency heat-maps confirmed depletion of Hi-C contact frequencies at all four *Hox* loci (*A, B, C* and *D*) in cells grown in 2i (Figure 3A, Supplementary Figure 2A).

**Figure 3:**
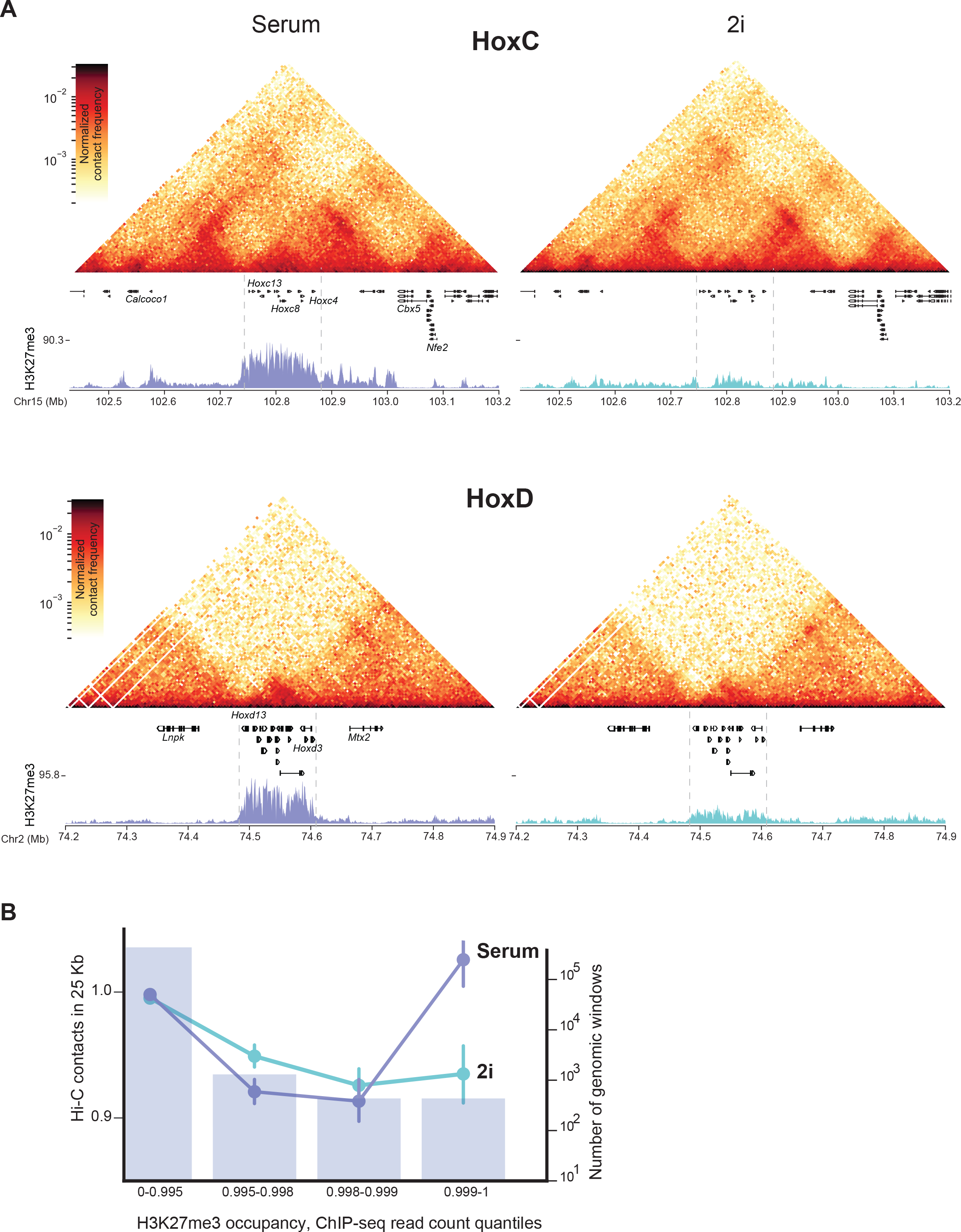
Loss of local chromatin interactions in 2i. A. Hi-C heatmaps (normalised contact frequencies) for cells grown in serum and 2i media for the *HoxC* (top) and *HoxD* (bottom) clusters along with gene annotations and H3K27me3 ChIP-seq profiles (Marks et al., 2012). Boundaries of the *Hox* clusters are marked with dashed lines. B. Quantification of local chromatin compaction measured as average number of observed/expected contacts in 25 kb sliding windows across different quantiles of H3K27me3 in serum and 2i media (coloured dots, mean ± 95% CI from bootstrapping, left *y* axis). Purple bars show the number of windows in each category (right *y* axis with log scale).

To investigate whether depletion of Hi-C contacts under 2i culture conditions is a global property of all polycomb targets, not just *Hox* loci, we divided the genome into sliding 25 kb windows. For each window we calculated the H3K27me3 ChIP-seq read density in either serum or 2i (Marks et al., 2012), and the number of Hi-C contacts. This showed (Figure 3B) that a high level of polycomb occupancy correlates with high local contact frequency in Hi-C in serum conditions, and that local Hi-C interactions are globally depleted in mESCs specifically at the genomic regions most enriched in H3K27me3. The same was observed when Hi-C contacts were assessed against RING1B occupancy (Illingworth et al., 2012) (Supplementary Figure 2B). In contrast, global analysis showed that local Hi-C interactions were similar between 2i and serum grown cells, as assessed by correlation of their insulation scores (Supplementary Figure 2C).

### Loss of polycomb-mediated long-range interactions in 2i-cultured mESCs

Polycomb has also been implicated in more long-range interactions using 4C, 5C and promoter-capture Hi-C (Bonev et al., 2017; Denholtz et al., 2013; Joshi et al., 2015; Kundu et al., 2017; Schoenfelder et al., 2015; Vieux-Rochas et al., 2015). Visual inspection of our Hi-C data within defined genomic windows in serum/LIF mESCs confirms that there are strong contacts between separate polycomb (H3K27me3) marked loci – for example between the *Skida1* and *Bmi1* loci separated by 650 kb on mouse chromosome 2 (Figure 4A), and between the *En2*, *Shh* and *Mnx1* loci across 1.3 Mb on chromosome 5 (Figure 4B). Consistent with the redistribution of H3K27me3 under 2i conditions, these long-range contacts are depleted or lost from cells in the ground state (Figure 4A and B). To analyse such interactions genome-wide, we used pile-up averaging of intra-chromosomal interactions between all CpG-islands (CGIs) either bound by PRC1 (RING1B), or not (Figure 4C, Supplementary Figure 3A). This showed reduced interactions under 2i conditions at CGIs specifically bound by RING1B, suggesting that the interactions disrupted under 2i are related to polycomb and not to general features of CGI promoters. Reduced interactions at polycomb sites were also confirmed by analysis of loops annotated in published Hi-C data from mESCs (Bonev et al., 2017). RING1B-associated loops across the genome display a clear depletion of interactions in 2i cells compared to those grown in serum (Figure 4D, Supplementary Figure 3B). In contrast, interactions between CTCF sites were not diminished and even seem enhanced in 2i. We also performed the same analysis on published Hi-C data from ICM/E3.5 embryo (Du et al., 2017; Ke et al., 2017; Zhang et al., 2018). While we cannot be sure of the polycomb distribution across the genome at this stage of embryogenesis in vivo, consistent with a DNA hypomethylated state, we observe high levels of enrichment for CTCF-associated loops in these datasets, but no enrichment at sites corresponding to RING1B-associated loops (Figure 4D).

**Figure 4:**
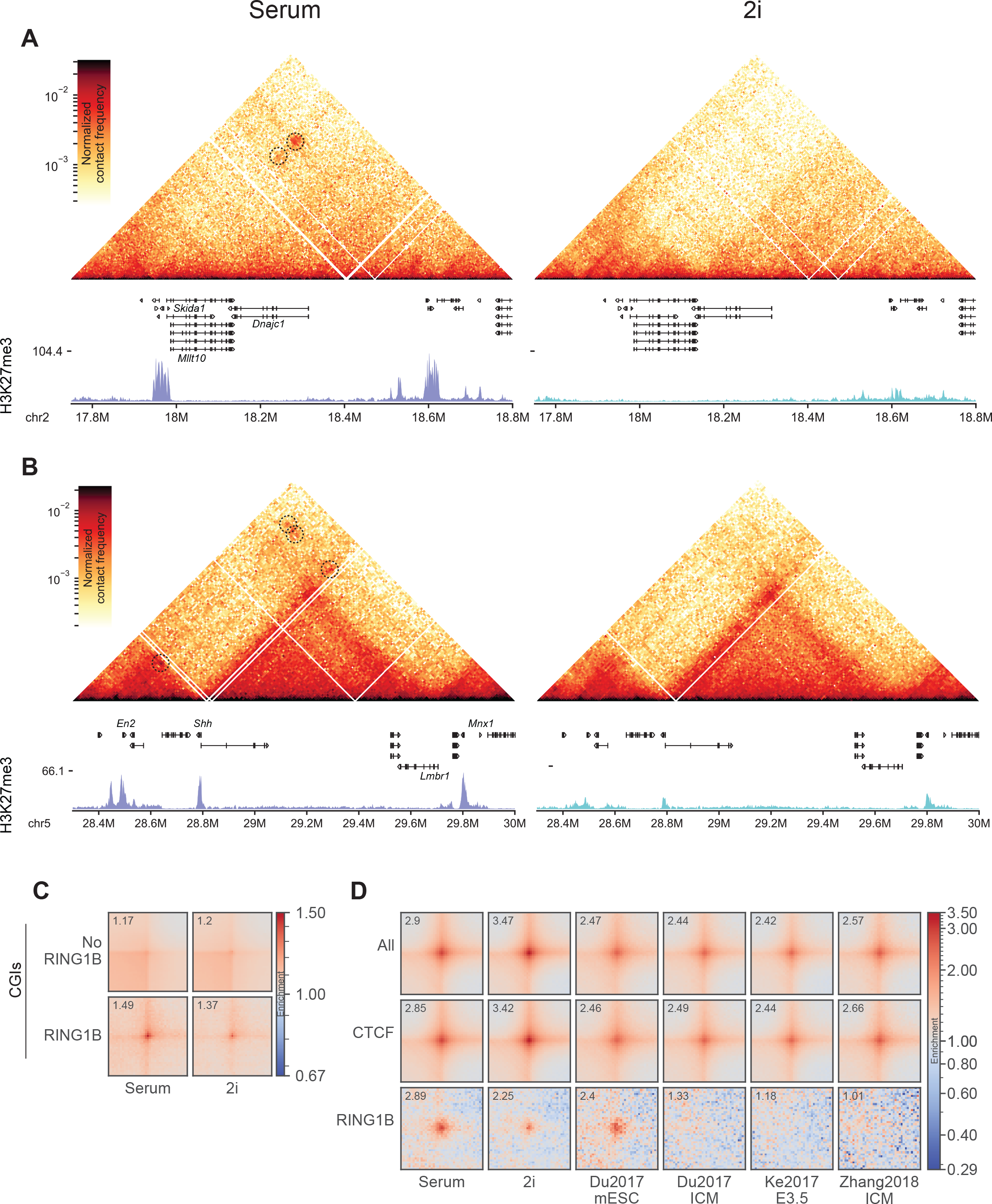
Loss of long-range chromatin interactions between polycomb loci in 2i. A. Example of a distal interaction between polycomb targets, *Bmi1* and *Skida1*, that is lost in 2i culture, along with gene annotations and H3K27me3 ChIP-seq profiles (Marks et al., 2012). Interactions in data from serum-cultured cells are highlighted with dashed circles. B. Same as A, but for interactions between *En2*, *Shh* and *Mnx1*. C. Averaged interactions (“pileups”) between CpG islands (CGIs) either occupied, or not, by RING1B in Hi-C data from serum and 2i cultured cells. Value of the centre pixel is shown in the top left corner of each heatmap. D. Pileups at loops called in mESC Hi-C data (Bonev et al., 2017), using our serum and 2i data, and compared to published Hi-C data from ICM (Du et al., 2017; Zhang et al., 2018/E3.5 embryos (Ke et al., 2017). Shown are all loops (All), those associated with CTCF peaks but not RING1B peaks (CTCF), and those associated with RING1B peaks (RING1B), but not CTCF peaks. Association is determined by the highest enriched pixel in the loop being within 5 kb of a ChIP-seq peak on both ends, while lack of a peak on at least one of the sides is treated as no association. Value of the centre pixel is shown in the top left corner of each heatmap.

### Preservation of DNA methylation in 2i prevents *HoxD* decompaction

The 3D chromatin re-organisation at polycomb targets we observe under 2i conditions could be a consequence of DNA hypomethylation-mediated polycomb redistribution or a reflection of the altered developmental potential of mESCs cultured in 2i relative to their serum counterparts. To distinguish between these two possibilities, we sought to uncouple the epigenetic transitions from the developmental changes in 2i cells.

DNA hypomethylation in 2i is thought to be the consequence of repression of *Dnmt3a*, *3b* and *Dnmt3l* by PRDM14 (Ficz et al., 2013; Yamaji et al., 2013). Therefore, we established a mESC line in which a high level of DNA methylation is maintained under 2i conditions. This was achieved utilising a DKO (*Dnmt3a*^−/−^, *3b*^−/−^) mESC cell line (3B3) in which DNA methylation is maintained with a Dnmt3B expressing transgene under the control of the CAG promoter (Jackson et al., 2004). In addition, we expressed the de novo methyltransferase co-factor Dnmt3L from a CAG promoter to create the mESC line 3B3L. Unlike the endogenous gene loci, the *Dnmt3b* and *Dnmt3l* transgenes in 3B3L cells are not repressed by PRDM14. HPLC confirmed that high CpG DNA methylation levels are retained in 3B3L cells cultured in 2i (Figure 5A). Consistent with the model where DNA methylation focuses polycomb targeting, ChIP-sequencing revealed that the maintenance of serum-level DNA methylation levels under 2i conditions in 3B3L cells also resulted in H3K27me3 being largely retained at polycomb target loci (Figure 5B and C).

**Figure 5:**
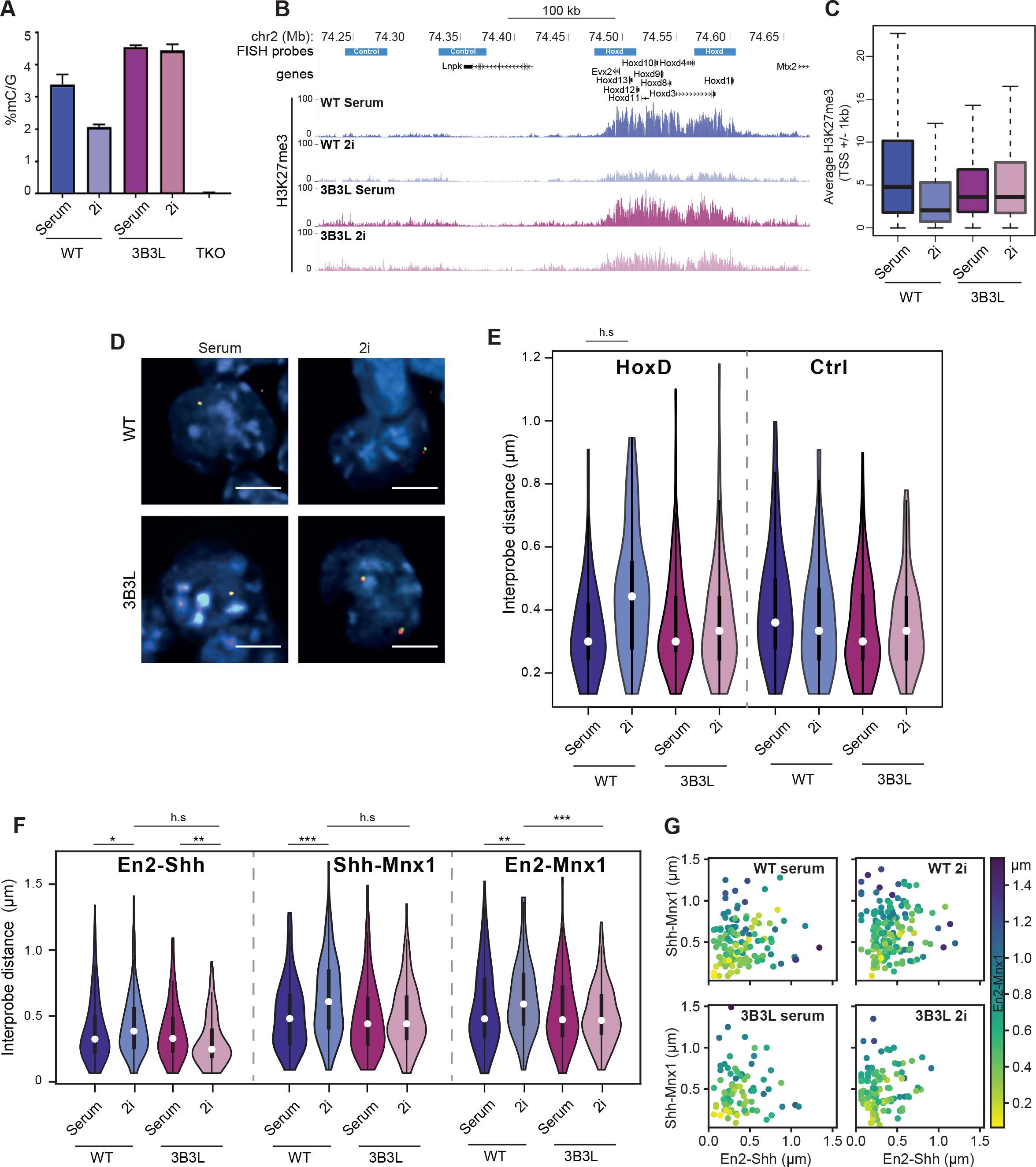
Restoration of the epigenetic and 3D landscape in 2i. A. DNA methylation measured by mass spectrometry showing global levels of methylated cytosine in WT and 3B3L cells under serum/LIF and 2i/LIF conditions (as well as negative control TKOs (Tsumura et al., 2006) which lack all the active DNMTs). Values represent % methylated cytosine normalised to total guanine. The mean of two technical replicates is shown, with error bars representing the standard deviation of the mean. B. UCSC genome browser screen shot at the *HoxD* locus showing H3K27me3 ChIP-seq in WT and 3B3L cells under serum/LIF and 2i/LIF conditions. Data for wild type cells is from Marks et al., (2012). Data are binned into 200 bp windows and normalised by total read count with reads from matching input samples subtracted. C. Boxplots representing average H3K27me3 signal on promoters (+/− 1 kb from TSS) for all promoters in WT or 3B3L cells under serum or 2i conditions D. Representative images of nuclei after FISH with probes for *HoxD* from WT and 3B3L cells grown in serum or 2i. Scale bars represent 10 μm. E. Violin plots showing distribution of inter-probe distances at the HoxD and a control (Ctrl) locus for WT J1 and 3B3L cells cultured in serum/LIF and 2i/LIF. h.s = p< 0.0001. Biological replicate for 3B3L cells, and data for 3A3L cells are in Supplementary Figure 4. F. Same as E, but for probes to En2, Shh, and Mnx1. G. Scatter plots showing individual measurements for data in F, with two distances shown along the axes and one (En2-Mnx1) colour-coded.

Consistent with the role of polycomb in mediating chromatin compaction, FISH revealed that the *HoxD* locus is retained in a compact chromatin confirmation when 3B3L cells are grown in 2i, contrasting with the decompaction seen at this locus when wild-type ESCs are grown in these culture conditions (Figure 5D). Inter-probe distances measured across *HoxD* were not significantly different between 3B3L cells grown in serum or 2i (Figure 5E and Supplementary Figure 4B). This result was also confirmed utilising a cell line in which *Dnmt3a* and *Dnmt3l* transgenes were exogenously expressed from a constitutive promoter (Supplementary Figure 4C).

Similarly, in contrast to the loss of long-range clustering between distant polycomb sites such as *En2*, *Shh* and *Mnx1* seen in WT ES cells in 2i (Figure 4B), inter-probe distances were not increased when 3B3L cells were cultured in 2i (Figure 5F) and the clustering of all three loci together was maintained (Figure 5G). This is consistent with the maintenance of H3K27me3 at these regions in 3B3L cells cultured under 2i conditions (Supplementary Figure 4A).

### The phenotype of 2i ESCs is driven by culture conditions not the epigenome or 3D chromatin organisation

Using 3B3L cells we are able to grow mESCs in 2i culture conditions but to largely maintain the epigenome and 3D genome organisation of ESCs grown in serum. To determine whether the phenotype of these cells is determined by the epigenome and 3D genome organisation or by the 2i condition and its impact on signalling, we first analysed features characteristic of the 2i naïve ground state of pluripotency.

3B3L cells still appear to exhibit hallmarks of the 2i ground state including upregulation of *Prdm14* (Figure 6A) and characteristic spheroid colony morphology. There was also uniform staining for ESRRB in 3B3L cells growing in 2i contrasting the heterogeneous staining seen in serum grown cells (Figure 6B). Serum and 2i mESCs have distinct transcriptional profiles (Marks et al., 2012). To determine whether the transcriptional profile of 3B3L cells in 2i more closely resembles that of mESCs with a similar epigenome and 3D organisation (serum ESCs), or that of mESCs grown under similar signalling blockade (2i), we compared RNA-seq data obtained from 3B3L and WT mESCs in 2i conditions. Principal component analysis showed that the 3B3L/2i transcriptome clusters with that of wild-type (J1) cells in the same condition, rather than with that of 3B3L cells grown in serum (Figure 6C). These results imply that the serum-like epigenome and 3D genome organisation of 3B3L cells growing in 2i conditions has little or no effect on the naïve pluripotency transcriptional state of these cells.

**Figure 6.**
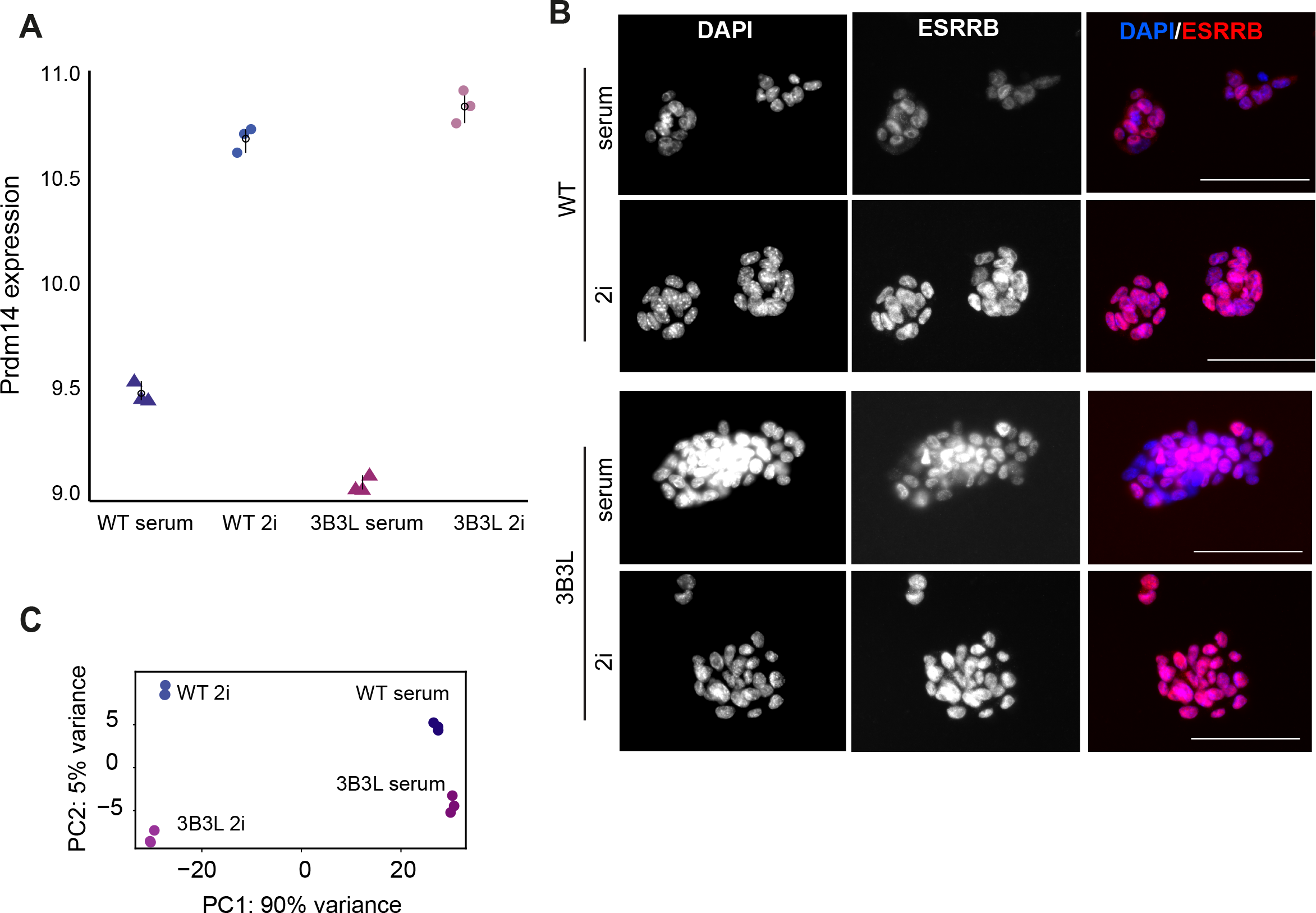
Analysis of the functional state of 3B3L mESCs in 2i. A. Regularized log (rlog) transformed expression value for Prdm14 in WT and 3B3L cells cultured in serum or 2i. Error bars show mean and bootstrapped 95% confidence intervals for each cell type and treatment group. Data are from 3 biological replicates B. ESRRB staining in 3B3L cells under 2i or serum conditions. Nuclei are counterstained with DAPI. Exposure times for the T×Red channel (ESRRB) were matched between conditions. Scale bars represent 100 μm. C. Principal component analysis (PCA) of the transcriptome (RNA-seq) of wild-type J1 and 3B3L mESCs cultured in serum or 2i. Data are from 3 biological replicates.

## Discussion

The observed patterns of DNA/histone modification profiled across the genome, and spatial genome organisation assayed by imaging or chromosome conformation capture assays, often correlate with patterns of gene regulation. However, experiments that determine whether there is a causal relationship between the epigenome, 3D genome and gene regulation are often lacking.

### DNA methylation impacts on 3D genome organisation via polycomb

As previously established by ourselves and others (Brinkman et al., 2012; Jermann et al., 2014; Marks et al., 2012; Reddington et al., 2013; Reddington et al., 2014), DNA methylation has a profound effect on the distribution of polycomb (H3K27 tri-methylation) across the mammalian genome, including in ESCs. This is likely to be a result of the specific targeting of both PRC2 and PRC1 to CpG islands (Farcas et al., 2012; Riising et al., 2014) and the generalised affinity of polycomb complexes for chromatin (Blackledge et al., 2015).

Here we have shown that the altered epigenome of 2i ESCs influences the 3D organisation of the genome; specifically, a loss of both local chromatin compaction at polycomb target loci (Figures 1 and 3) and long-range polycomb-mediated chromatin contacts (Figures 4 and 5). We also show that the loss of chromatin compaction at polycomb target loci, such as *Hox* loci, in naïve pluripotency reflects their chromatin conformation in vivo in hypomethylated preimplantation blastocysts (Figure 2).

In contrast to polycomb target loci, chromatin compaction at a control non-polycomb target locus was not significantly different between mESCs grown in serum vs 2i conditions (Figure 1). This suggests that chromatin decompaction is not a result of a global alteration in the 3D chromatin organisation of naïve 2i cells. It also demonstrates that the significant loss of DNA methylation across a genomic region has no detectable effect of chromatin compaction, as assayed at a cytological level. This is consistent with the finding that chromatin compaction as assayed by nuclease sensitivity and sucrose gradient sedimentation in mammalian cells is also not affected by the loss of DNA methylation (Gilbert et al., 2007).

### The epigenome and 3D genome do not affect the naive pluripotency functional state

By manipulating the epigenome (DNA methylation and H3K27me3 distribution) of mESCs grown in 2i conditions we have been able to demonstrate that changes in 3D genome organisation that occur as ESCs transition between primed and naïve pluripotency, are a downstream consequence of the shifting epigenome. Constitutive expression of de novo DNA methyltransferases during the conversion to 2i conditions largely prevents the changes to DNA methylation and polycomb targeting normally seen for wild-type mESCs in these culture conditions (Figure 5). This is then reflected in 3D genome organisation – in 3B3L mESCs cultured in 2i, *Hox* loci retain their local chromatin compaction and long-range clustering of polycomb sites is preserved (Figure 5).

However, this ‘serum-like’ epigenome and 3D genome organisation that we have imposed on ESCs growing in 2i does not detectably affect the transcriptional state of the ESCs. They maintain their high and homogeneous expression of pluripotency markers and the transcriptome of these cells resembles that of the ground state (2i), not that of primed (serum) ESCs (Figure 6). This is consistent with the observation that the decrease of H3K27me3 at gene promoters is not generally associated with transcription activation of these loci under 2i conditions (Galonska et al., 2015; Marks et al., 2012; van Mierlo et al., 2019). We propose that the transcriptional network driven by the defined 2i signalling environment predominates over any instructive information in the epigenome or 3D genome.

Our data demonstrate that 3D genome organisation is an emergent property of the epigenome and that the consequences of perturbing one part of the epigenome (DNA methylation) cannot be considered in isolation. Rather the impact of one epigenetic system on other epigenetic systems (e.g. polycomb) and the related changes in 3D genome organisation must be considered together. Our findings also caution against over-interpreting the functional significance of the epigenome and 3D genome organisation – at least in ESCs. It will now be interesting to establish how DNA methylation and polycomb become increasingly functionally important for gene regulation as development progresses (Greenberg et al., 2017).

## Acknowledgments

We are grateful to Nezar Abdennur, Anton Goloborodko and Maxim Imakaev for advice on Hi-C data analysis, and to Sergey Venev for prompt help with loop annotation. We thank other Bickmore and Meehan lab members for discussions. KMcL was funded by a PhD studentship from the UK Medical Research Council (MRC). IMF was funded by a PhD studentship from the Darwin Trust. Work in the RMM lab is funded by an MRC University Unit grant MC_PC_U127574433. Work in the SP lab was funded by the BBSRC and BHF. IRA is funded by an MRC University Unit grant MC_PC_U127580973. Work in the WAB lab is funded by an MRC University Unit grant MC_UU_00007/2.

## Author contributions

The study was conceived and designed by RRM and WAB. FISH analysis of *Hox* loci was performed and analysed by KMcL. FISH analysis of *En2*, *Shh* and *Mnx1* was performed by IW. Hi-C was performed by IMF and KMcL and analysed by IMF. HKM, RS and SP created and characterised stable cell lines 3A3L and 3B3L, performed ChIP-seq, prepared samples for RNA-seq and assisted with experimental design and protocols. I.R.A. provided blastocysts for FISH. KMcL, IMF, GRG, and WAB prepared figures. RNA-Seq analysis was performed by GG. JPT and RSI performed computational genomic analysis. KMcL, RM, WAB and IMF wrote the manuscript with input from all authors.

## STAR Methods

### EXPERIMENTAL MODELS

Mouse embryonic stem cell lines used in this study are: E14, WT (clone 36), *Ring1B*^−/−^, *Eed*^−/−^ (Eskeland et al., 2010) and WT J1. 3B3L/3A3L cells are DKO (*Dnmt3a*^−/−^, *3b*^−/−^) mESC lines where DNA methylation is maintained with a Dnmt3b/ or Dnmt3a expressing transgene under the control of the CAG promoter (3B3/3A3) (Jackson et al., 2004; Okano et al., 1999) and which were transfected with pCAGGS-*Dnmt3l-Flag*-IRES-Blasticidin-polyA, and selected by blasticidin (5 μg/ml) to obtain cell lines with stable expression of *Dnmt3l*.

### EXPERIMENTAL METHODS

#### Animals

Animal were maintained in accordance with institutional guidelines and national regulations. Animal experiments were performed under the authority of UK Home Office project licence PPL60/4424 following ethical review by the University of Edinburgh Animal Welfare and Ethical Review Body.

#### Cell culture

mESCs were maintained at 37°C with 5% CO2 and passaged every 2-3 days. Serum cells were maintained in either DMEM (in the case of J1-derived lines) or GMEM (in the case of E14-derived lines) (both Gibco) supplemented with 15% foetal calf serum, 0.1 mM nonessential amino acids (SIGMA), 1 mM sodium Pyruvate (Sigma) 1% Penicillin/Streptomycin, 2 mM L-glutamine, 0.1 mM β-mercaptoethanol (Thermo Fisher), and ESGRO LIF (Millipore) at 1000 U/mL. Cells were either grown on 0.2% gelatin (Sigma) (E14 cells, 3B3L cells) or on mitomyin C-inactivated SNLP feeder cells in the case of serum culture J1/clone36-derived cells. 2i culture conditions include 50% DMEM/F12 (Gibco), 50% Neurobasal media (Gibco), 0.5% N2 supplement, 1% B27 & RA (Gibco), 7.5% BSA (Gibco), 1% Penicilllin/Streptomycin, 2 mM L-glutamine, 0.15 mM monothioglycerol (Sigma), 1000 U/ml ESGRO LIF (Millipore), 1 μM PD0325901 (MEK inhibitor, Stemgent) and 3 μM CHIR99021 (GSK3 inhibitor, Stemgent). mESCs were passaged every 2-3 days using trypsin/EDTA (Sigma). 2i conversions were carried out for 14 days. To deplete feeder-dependent mESCs of their feeders for analysis/2i-conversion, the culture was plated 3x for 20 mins, in which time the feeders stick to the tissue culture dish and mESCs do not.

#### FISH

One million mESCs were plated onto gelatinised slides for 4 h. Cells were fixed in 4% paraformaldehyde (pFA) for 10 min, permeabilized in 0.5% Triton X-100 for 10 min, air-dried and stored at −80 °C. Slides were incubated with 100 ug/ml RNaseA in 2 × SSC for 1 h, washed in 2 × SSC and dehydrated through an alcohol series. Slides were then denatured in 70% formamide/2xSSC at 80°C for 30 min. Fosmid clones (Supplementary Table 1) were prepared and labelled with digoxigenin-11-dUTP or with biotin-16-dUTP as previously described (Morey et al., 2007), or directly labelled with fluorescent nucleotides (approximately 80 ng of biotin- and digoxigenin-labelled fosmid probes were used per slide, 1 mM ChromaTide Alexa 594-5-dUTP (Invitrogen) or 2.5 μl of 1 mM 5(6)-Carboxyrhodamine Green dUTP (Enzo).

Approximately 150 ng of labelled fosmid probes were used per slide, together with 8 μg of mouse Cot1 DNA (Invitrogen) and 5 μg sonicated salmon sperm DNA. Probes were denatured at 80°C for 5 min, preannealed for 15 min at 37°C and hybridized to the denatured slides overnight (o/n). The following day, the slides were washed in 2x SSC followed by 0.1x SSC, and stained in DAPI prior to imaging.

For FISH on blastocysts, an adaptation of previously described protocols was used (Flyamer et al., 2017; Probst et al., 2007). Briefly, 20 female C57BL/6 mice were superovulated and mated with C57BL/6 males, and blastocysts isolated at E3.5 by flushing the uterine horns with FHM media. Blastocysts with visible blastocoels were fixed in 4% pFA and their zona pellucidae removed using Acidic Tyrode’s. The blastocysts were permeabilised in 0.2% Triton-X 100 in PBS. Fixed samples were embedded in fibrin clots to attach the blastocysts to slides. Post-fixation was carried out in 2% pFA/ PBS for 30 min. Finally, the slide was rinsed 3x in PBS and stored in PBS at 4°C. FISH was carried out using directly labelled probes described above, with some modifications. The slides were denatured for 45 min.

#### Image capture

Images were captured using a Hamamatsu Orca AG CCD camera (Hamamatsu Photonics (UK) Ltd, Welwyn Garden City, UK) and a Zeiss Axioplan II epifluorescence microscope with Plan-neofluar objectives, a 100W Hg source (Carl Zeiss, Welwyn Garden City, UK) and Chroma #83000 triple band pass filter set (Chroma Technology Corp., Rockingham, VT) with the excitation filters installed in a motorised filter wheel (Prior Scientific Instruments, Cambridge, UK). A piezoelectrically driven objective mount (PIFOC model P-721, Physik Instrumente GmbH & Co, Karlsruhe) was used to control movement in the z dimension (with 0.2 μm step).

#### Immunocytochemistry

mESCs grown on glass coverslips coated with gelatin were fixed with 4% PFA for 20 mins, blocked in 10% donkey serum (Sigma) in 0.1% Triton X-100 for 1 h and incubated o/n with primary antibody detecting ESRRB (Perseus Proteomics, PP-H6705-00) at a 1:500 dilution at 4°C. The following day, samples were incubated with Donkey anti mouse Alexafluor 555 (Cat: A-31570, Thermo Fisher) at room temperature for 1 h. Nuclei were counterstained with DAPI. Imaging was carried out using a Zeiss Axioscope 2 microscope.

#### H3K27me3 ChIP-seq

Chromatin prepared from formaldehyde fixed 3B3-3l cells cultured in serum or 2i was fragmented (Covaris sonicator) to a mean fragment size of 200bp. Approximately 5×10^6^ cell equivalents were used for each immunoprecipitation. ChIP was performed using antibody towards H3K27Me3 (Millipore) and Protein G Dynabeads (Thermo Fisher) were used to obtain antibody bound chromatin. Following immunoprecipitation, beads were washed once in X-ChIP wash buffer (150mM NaCl; 10mM Tris pH8; 2mM EDTA; 1% NP40; 0.1% sodium deoxycholate w/v), and once in LiCl wash buffer (100mM Tris pH7.5; 500mM LiCl; 1% NP40; 1% sodium deoxycholate) for 10 min at 4°C each wash. DNA was then reverse crosslinked and eluted from the beads by incubation in elution buffer (1% SDS, 0.1M NaHCO_3_) followed by treatment with RNAse and proteinase K before purification using a Qiagen minelute kit (Qiagen) as per manufacturer’s instructions and eluting the DNA in 11μl EB buffer from the kit. Finally, DNA was quantified using a Qubit HS DNA quantification kit (Thermo Fisher) and 1ng DNA was then used to prepare sequencing libraries for Ion Torrent sequencing using the Ion XpressPlus Fragment Library Kit (Thermo Fisher). The DNA was end repaired, purified and ligated to Ion-compatible barcoded adapters (Ion Xpress™ Barcode Adapters 1–96: (Thermo Fisher) followed by nick-repair to complete the linkage between adapters and DNA inserts. The adapter-ligated library was then amplified (10 cycles) and size-selected using two rounds of AMPure XP bead (Beckman Coulter) capture to size‐select fragments approximately 100–250bp in length. Samples were pooled at a 1:1 ratio and sequenced on an Ion Proton P1 microwell chip (Thermo Fisher).

Mapping and data normalization were carried out as described previously (Thomson et al., 2015). In short, reads were mapped to the reference genome using the Torrent TMAP software. The data was then binned into 200bp windows across the genome and normalised by total read count. Raw sequencing datasets from published WT E14 mESCs in both serum and 2i were processed in a similar manner (Marks et al., 2012): NCBI GSE23943.

#### RNA-seq

RNA was extracted from snap frozen mESC pellets, 3 biological replicates per cell line, using an RNeasy kit (Qiagen). RNA was quantified by nanodrop and DNA was removed by treatment with Turbo DNA-free reagents (AM1907, Ambion) according to the manufacturer’s protocol. Total RNA samples were quantified using the Qubit 2.0 Fluorometer (Thermo Fisher Scientific Inc, Q32866) and the Qubit RNA HS assay kit (Q33855). RNA integrity was assessed using the Agilent 2100 Bioanlyser System (Agilent Technologies Inc, GS2938B) and Agilent RNA 6000 Nano kit (5067-1511).

Sequencing libraries were prepared from 500 ng of each total-RNA sample using the TruSeq Stranded mRNA Library Kit (Illumina Inc, 20020594) according to the provided protocol. Poly-A mRNAs were purified using poly-T oligo attached magnetic beads, and fragmented using divalent cations under elevated temperature and primed with random hexamers. Primed RNA fragments were reverse transcribed into first strand cDNA using reverse transcriptase and random primers. RNA templates were removed and a replacement strand synthesised incorporating dUTP in place of dTTP to generate ds cDNA. The incorporation of dUTP in second strand synthesis quenches the second strand during amplification as the polymerase used in the assay is not incorporated past this nucleotide. AMPure XP beads (Beckman Coulter, A63881) were then used to separate the ds cDNA from the second strand reaction mix, providing blunt-ended cDNA. A single ’A’ nucleotide was added to the 3’ ends of the blunt fragments to prevent them from ligating to one another during the subsequent adapter ligation reaction, and a corresponding single ’T’ nucleotide on the 3’ end of the adapter provided a complementary overhang for ligating the adapter to the fragment. Multiple indexing adapters were then ligated to the ends of the ds cDNA to prepare them for hybridisation onto a flow cell, before 12 cycles of PCR were used to selectively enrich those DNA fragments that had adapter molecules on both ends and amplify the amount of DNA in the library suitable for sequencing. After amplification libraries were purified using AMPure XP beads.

Libraries were quantified by fluorometry using the Qubit dsDNA HS assay and assessed for quality and fragment size using the Agilent Bioanalyser with the DNA HS Kit (5067-4626). Sequencing was performed using the NextSeq 500/550 High-Output v2 (150 cycle) Kit (FC-404-2002) on the NextSeq 550 platform (Illumina Inc, SY-415-1002). Twenty four libraries were combined in two equimolar pools of 12 based on the library quantification results and each pool was run across a single High-Output Flow Cell. Sequencing was performed at the Wellcome Trust Clinical Research Facility (WTCRF; Edinburgh).

#### DNA methylation by mass spectrometry

DNA was extracted from frozen cell pellets by standard phenol:chloroform extraction and ethanol purification. 2.5 μg DNA in 50 μL final volume was then hydrolysed by incubation at 95 °C for 10 mins and incubated overnight at 37 °C with 5 μL T7 DNA polymerase reaction buffer and 1 μL 10 U/μL T7 DNA polymerase (Thermo Fisher). The reaction was heat inactivated at 75°C for 10 mins. The sample was then centrifuged at 12,000×g at r.t. for 45 mins.

To carry out DNA hydrolysis, 2.5μg of DNA was made up to 44μl in mass spectrometry grade water (Chromasolv, Sigma) and incubated at 95°C, 10 mins. 5μl T7 DNA polymerase reaction buffer and 1μl 10U/μl T7 DNA polymerase (Thermo Fisher) were added and the samples incubated o/n at 37°C. The reaction was heat inactivated at 75°C for 10 mins. The sample was then centrifuged at 12,000 ×G at r.t. for 45 mins.

The hydrolysed DNA was then extracted in 5:3:2 methanol:acetonitrile:sample, and centrifuged at 12,000 ×g for 5 mins, the upper 90 μL were taken and the organic solvent removed using a vacuum centrifuge. Analytes were resuspended in 30 μL mass spectrometry grade water and 10 μL injected onto a 30x 1mm HyperCarb column (VWR). A gradient of 0-90% B was run over 4 mins, where B is acetonitrile and A is 20 mM ammonium carbonate. Mass spectra were acquired in negative mode on a Thermo Q Exactive, scanning from 300 to 350 m/z at resolution 70k. AGC target was set to 1x 106 and maximum ion time 100ms. Data were analysed using AssayR (Wills et al., 2017).

#### In situ Hi-C

Hi-C was performed largely as described (Rao et al., 2014) with minor modifications. Briefly, 2-5×10^6^ mESCs were crosslinked in 1% formaldehyde for 10 mins, snap-frozen and stored at −80 °C. After permeabilization in lysis buffer (0.2% Igepal, 10 mM Tris-HCl pH 8.0, 10 mM NaCl, 1x Halt Protease inhibitor cocktail) nuclei were isolated in 0.3% SDS in NEBuffer 3 at 62°C for 10 min. SDS was quenched with 1% Triton X-100 at 37°C for 1 h, then the nuclei were pelleted and resuspended in 250 μl DpnII buffer with 600 U DpnII. After digestion o/n, 200 more units were added for 2 h. Then the ends were filled-in using Klenow, d(G/C/T)TPs and biotin-14-dATP for 1.5 h at 37°C. After ligation at room temperature for 4 h the nuclei were spun down, resuspended in 200 μl mQ and digested with proteinase K for 30 min at 55°C in presence of 1% SDS. Cross-links were reversed at 65°C o/n after addition of NaCl to a final concentration of 1.85 M. After ethanol precipitation and a 70-80% ethanol wash, DNA was resuspended in 500 μl of sonication buffer (50 mM Tris pH 8.0, 0.1% SDS, 10 mM EDTA), incubated on ice for 15 min and then sheared using a probe sonicator to fragment sizes of 200-700 bp. DNA was then concentrated on Amicon filter units, bound to MyOne T1 Streptavidin beads and used for Illumina library preparation. Small aliquots were taken before and after DpnII treatment, and before sonication to confirm efficient DNA digestion and ligation by running them on 1% agarose gel. Samples were first test-sequenced on NextSeq 550 (WTCRF, Edinburgh) to check library quality, and then selected libraries were sequenced at greater depth on HiSeq 4000 (BGI-Hongkong).

### QUANTIFICATION AND STATISTICAL ANALYSIS

#### FISH image analysis

Volocity software (PerkinElmer) was used to capture, process and analyse the images. Images were deconvolved using the Restoration module, using the constrained iterative algorithm. Image analysis was carried out using the Quantitation module. For analysis of data from ESCs, each data set consisted of 70-155 measurements. For analysis of Hox probes in blastocysts, 686 alleles from 14 embryos were analysed. Control inter-probe distances were measured from 100 alleles from 2 blastocysts. The statistical analysis of inter-probe distance distributions was determined using a Mann-Whitney U Test. Mean inter-probe distances for all FISH data are shown in Supplementary Table 2 and p values are listed in Supplementary Table 3.

#### Hi-C data analysis

Reads were processed using distiller (https://github.com/mirnylab/distiller-nf) on the high-performance computing cluster of the University of Edinburgh (Eddie). Mapping was performed to the mm9 genome build. Hi-C pairs with exactly matching coordinates were removed as PCR or optical duplicates (pcr_dups_max_mismatch_bp: 0). The output statistics information and Cooler files (https://github.com/mirnylab/cooler) were used in downstream analyses. 1000 bp resolution Cooler files were used to create multi-resolution files for visualization in HiGlass.

For pileup analysis, we took all regions of interest in the Hi-C maps, e.g. all *cis* interactions between CGIs bound or not bound by RING1B (Illingworth et al., 2015)), and averaged a 15×15 window centred on them at 5 Kb resolution (10 Kb for the Du et al data, and comparison with it). For each of those windows, 10 control windows were obtained by randomly shifting the position of the original window between 100 kb and 1 Mb along the diagonal of the map. All control windows were averaged and used to normalise the pile-up to remove the distance-dependency of interactions. Then, the final matrix was divided by the average value of its top-left and bottom-right 3×3 corners, to normalize the local background. Values of enrichment are the enrichment of interactions in the centre pixel of the matrix, after all described normalization procedures. The code used to perform this analysis is available here: https://github.com/Phlya/coolpuppy.

Since our own Hi-C data was not deep enough to call loops with high quality, we chose instead to take advantage of very deeply sequenced published data from mESCs (Bonev et al., 2017). We used *cooltools call-dots* reimplementation of the HiCCUPS algorithm (Rao et al., 2014) from *dekkerlab/shrink-donut-dotfinder* (commit 377106e). This was applied with default settings (except for lower FDR threshold of 0.1) to reanalysed mapq≥30 filtered mESC Hi-C data at 5 kb, 10 kb and 25 kb resolution to find areas of local enrichment of interactions between loci up to 20 Mb away. Calls from different resolutions were combined using a custom script following the HiCCUPS merging procedure. Annotated dots were then filtered by intersecting with published CTCF peaks (Bonev et al., 2017), and/or RING1B peaks (Illingworth et al., 2015) using *bedtools pairtobed* after widening the peaks using *bedtools slop*.

For local interaction density analysis, we used 5 kb resolution data and sliding 25 kb windows with a 5 kb step. For each window we determined average observed/expected number of interactions (excluding the first two diagonals, so we averaged 6 pixels per window). If at least 20% of the pixels in the window were missing (NaN), we did not consider it (i.e. >1 pixel, however since missing values come from masking whole genomic bins during balancing, effectively having one masked bin removed the window from analysis). Then these data were combined with the read coverage in the same windows from ChIP-seq experiments (H3K27me3 from (Marks et al., 2012), RING1B from (Illingworth et al., 2015)). Binning of the windows into groups was performed based in quantiles of ChIP-seq values, and mean (±95% confidence interval obtained by bootstrapping) was plotted using *seaborn python* package, together with the total number of windows considered in the analysis after all filtering.

For insulation score analysis, we applied *cooltools diamond-insulation* with window size of 100 kb. We then discarded any invalid bins, and performed pairwise Pearson correlation analysis between individual samples to assess the similarity between the samples.

The GEO accession for Hi-C data is GSE124342. The data can be visualised through the following link http://higlass.io/app/?config=Gi42-PrxQYCGeTDuGEPWLA

#### RNA-seq analysis

RNA-Seq data can be accessed from the Gene Expression Omnibus (https://www.ncbi.nlm.nih.gov/geo) using series accession number GSE121171.

mRNA abundance was quantified using Sailfish (version: 0.9.2,-l ISR) against mm10 transcript models as defined by RefSeq. The R package tximport was used to import and summarize transcript-level estimates for gene-level analysis. The regularized log transformation (rlog, R Package DESeq2) was applied to minimizes differences between samples for rows with small counts, and which normalizes with respect to library size. To visualize sample-to-sample distances a principal component analysis (PCA) was performed using the rlog values.

