## Supplementary information for "DNA methylation directs polycomb-dependent 3D genome reorganisation in naïve pluripotency"

### Supplemental Data

**Supplementary Table 1** Details of FISH probes

Genome co-ordinates are given using the mm9 assembly of the mouse genome

| Locus | Whitehead Name | Coordinates (mm9) | Size (bp) |
|---|---|---|---|
| *Hoxd3* | WI1-121N10 | Chr2: 74,566,983 – 74,605,438 | 38,455 |
| *Hoxd13* | WI1-469P2 | Chr2: 74,474,157 -74,513,003 | 38,846 |
| *GCR* | WI1-2157A11 | Chr2: 74,242,615 -74,282,044 | 39,429 |
| *Lnp* | WI1-482L15 | Chr2: 74,329,582 -74,372,986 | 43,404 |
| *Hoxb1* | WI1-2671L18 | Chr11: 96,201,164- 96,242,956 | 41,793 |
| *Hoxb13* | WI1-1356F15 | Chr11: 96,060,900-96,099,631 | 38,732 |
| *Hoxc4* | WI1-0991J24 | Chr15: 103,018,285 -103,057,123 | 38,838 |
| *Hoxc13* | WI1-1176M4 | Chr15: 102,910,943 - 102,949,345 | 38,403 |
| *En2* | WI1-2728F4 | Chr5: 28,477,913 – 28,517,563 | 39,650 |
| *Shh* | WI1-574O18 | Chr5: 28,754,458 – 28,795,879 | 41,421 |
| *Mnx1* | WI1-1204B6 | Chr5: 29,791,124 – 29,827,491 | 36,367 |

**Supplementary Table 2.** Median inter-probe distances for each FISH probe set in all cell lines and conditions. Data from replicate experiments are indicated

| Cell line/condition | Probe pair | Median interprobe distance (µm) |
|---|---|---|
| **Figure 1 & Figure S1** | | |
| WT (clone36) Serum | *Hoxd3-Hoxd13* | Rep1: 0.276, Rep2: 0.317 |
| WT (clone36) 2i | *Hoxd3-Hoxd13* | Rep1: 0.424, Rep2: 0.423 |
| WT (clone36) Serum | *GCR-Lnp* | Rep1: 0.334, Rep2: 0.36 |
| WT (clone36) 2i | *GCR-Lnp* | Rep1: 0.36, Rep2: 0.422 |
| *Ring1B$^{-/-}$* Serum | *Hoxd3-Hoxd13* | Rep1: 0.379, Rep2: 0.443 |
| *Ring1B$^{-/-}$* 2i | *Hoxd3-Hoxd13* | Rep1: 0.483, Rep2: 0.459 |
| *Ring1B$^{-/-}$* Serum | *GCR-Lnp* | Rep1: 0.3, Rep2: 0.36 |
| *Ring1B$^{-/-}$* 2i | *GCR-Lnp* | Rep1: 0.3, Rep2: 0.334 |
| *Eed$^{-/-}$* Serum | *Hoxd3-Hoxd13* | Rep1: 0.483, Rep2: 0.481 |
| *Eed$^{-/-}$* 2i | *Hoxd3-Hoxd13* | Rep1: 0.39, Rep2: 0.469 |
| *Eed$^{-/-}$* Serum | *GCR-Lnp* | Rep1: 0.334, Rep2: 0.334 |
| *Eed$^{-/-}$* 2i | *GCR-Lnp* | Rep1: 0.36, Rep2: 0.334 |
| E14 Serum | *Hoxb1-Hoxb13* | 0.276 |
| E14 2i | *Hoxb1-Hoxb13* | 0.347 |
| E14 Serum | *Hoxc4-Hoxc13* | 0.3 |
| E14 2i | *Hoxc4-Hoxc13* | 0.36 |
| **Figure 2** | | |
| Blastocysts | *Hoxd3-Hoxd13* | 0.422 |
| Blastocysts | *GCR-Lnp* | 0.334 |
| **Figure 5 & S4** | | |
| WT J1 Serum | *Hoxd3-Hoxd13* | 0.3 |

| | | |
|---|---|---|
| WT J1 2i | *Hoxd3-Hoxd13* | 0.443 |
| WT J1 Serum | *GCR-Lnp* | 0.36 |
| WT J1 2i | *GCR-Lnp* | 0.334 |
| 3B3L Serum | *Hoxd3-Hoxd13* | Rep1: 0.3, Rep2: 0.3 |
| 3B3L 2i | *Hoxd3-Hoxd13* | Rep1: 0.334, Rep2: 0.334 |
| 3B3L Serum | *GCR-Lnp* | 0.3 |
| 3B3L 2i | *GCR-Lnp* | 0.334 |
| 3A3L Serum | *Hoxd3-Hoxd13* | 0.3 |
| 3A3L 2i | *Hoxd3-Hoxd13* | 0.36 |
| WT J1 Serum | *En2-Shh* | 0.324 |
| WT J1 2i | *En2-Shh* | 0.385 |
| WT J1 Serum | *Shh-Mnx1* | 0.48 |
| WT J1 2i | *Shh-Mnx1* | 0.608 |
| WT J1 Serum | *En2-Mnx1* | 0.478 |
| WT J1 2i | *En2-Mnx1* | 0.59 |
| 3B3L Serum | *En2-Shh* | 0.329 |
| 3B3L 2i | *En2-Shh* | 0.247 |
| 3B3L Serum | *Shh-Mnx1* | 0.44 |
| 3B3L 2i | *Shh-Mnx1* | 0.44 |
| 3B3L Serum | *En2-Mnx1* | 0.471 |
| 3B3L 2i | *En2-Mnx1* | 0.466 |

**Supplementary Table 3** Probability values calculated by Mann-Whitney U tests comparing inter-probe distances between two populations. Inter-probe distances of *HoxD*, *Lnp-GCR* (Ctrl), *HoxC*, and *HoxB* probes in different cell types and conditions are shown

| Genotype/ Condition/ Probes Sample 1 | Genotype/ Condition/ Probes Sample 2 | P-value (Mann-Whitney U Test) |
|---|---|---|
| **Figure 1 & Figure S1** | | |
| WT (clone36) Serum HoxD | WT (clone36) 2i *HoxD* | Rep1: <0.0001, Rep2: 0.0003 |
| WT (clone36) Serum HoxD | Ring1B $^{-/-}$ Serum *HoxD* | Rep1: <0.0001, Rep2: <0.0001 |
| WT (clone36) Serum HoxD | Eed $^{-/-}$ Serum *HoxD* | Rep1: <0.0001, Rep2: <0.0001 |
| WT (clone36) Serum HoxD | Ring1B $^{-/-}$ 2i *HoxD* | Rep1: <0.0001, Rep2: <0.0001 |
| WT (clone36) Serum HoxD | Eed $^{-/-}$ 2i *HoxD* | Rep1: <0.0001, Rep2: <0.0001 |
| WT (clone36) Serum Ctrl | WT (clone36) 2i Ctrl | Rep1: 0.4215, Rep2: 0.2564 |
| WT (clone36) Serum Ctrl | Ring1B $^{-/-}$ Serum Ctrl | Rep1: 0.1352, Rep2: 0.5583 |
| WT (clone36) Serum Ctrl | Eed $^{-/-}$ Serum Ctrl | Rep1: 0.7539, Rep2: 0.4776 |
| WT (clone36) Serum Ctrl | Ring1B $^{-/-}$ 2i Ctrl | Rep1: 0.0797, Rep2: 0.1836 |
| WT (clone36) Serum Ctrl | Eed $^{-/-}$ 2i Ctrl | Rep1: 0.5062, Rep2: 0.0865 |
| E14 Serum *HoxB* | E14 2i *HoxB* | 0.0334 |
| E14 Serum *HoxC* | E14 2i *HoxC* | 0.0024 |
| **Figure 2** | | |
| WT serum *HoxD* | Blastocysts *HoxD* (all) | <0.0001 |
| WT serum *HoxD* | Blastocyst 1 *HoxD* | 0.0050 |
| WT serum *HoxD* | Blastocyst 2 *HoxD* | 0.0170 |
| WT serum *HoxD* | Blastocyst 3 *HoxD* | 0.0168 |
| WT serum *HoxD* | Blastocyst 4 *HoxD* | 0.0043 |
| WT serum *HoxD* | Blastocyst 5 *HoxD* | <0.0001 |
| WT serum *HoxD* | Blastocyst 6 *HoxD* | <0.0001 |

| | | |
|---|---|---|
| WT serum *HoxD* | Blastocyst 7 *HoxD* | 0.0068 |
| WT serum *HoxD* | Blastocyst 8 *HoxD* | 0.0002 |
| WT serum *HoxD* | Blastocyst 9 *HoxD* | 0.0019 |
| WT serum *HoxD* | Blastocyst 10 *HoxD* | 0.1389 |
| WT serum *HoxD* | Blastocyst 11 *HoxD* | 0.0537 |
| WT serum *HoxD* | Blastocyst 12 *HoxD* | 0.0047 |
| WT serum *HoxD* | Blastocyst 13 *HoxD* | 0.0237 |
| WT 2i *HoxD* | Blastocysts *HoxD* (all) | 0.0879 |
| WT Serum Ctrl | Blastocysts Ctrl | 0.6855 |
| WT 2i Ctrl | Blastocysts Ctrl | 0.2017 |
| **Figure 5 and Figure S4** | | |
| WT J1 Serum *HoxD* | WT J1 2i *HoxD* | Rep1: <0.0001,Rep2: |
| WT J1 Serum *HoxD* | 3B3L Serum *HoxD* | Rep1: 0.2027, Rep2: |
| WT J1 Serum *HoxD* | 3B3L 2i *HoxD* | Rep1: 0.2790, Rep2: |
| 3B3L Serum *HoxD* | 3B3L 2i *HoxD* | Rep1: 0.8658 Rep2: |
| WT J1 Serum *HoxD* | 3A3L Serum *HoxD* | 0.7219 |
| WT J1 Serum *HoxD* | 3A3L 2i *HoxD* | 0.1128 |
| 3A3L Serum *HoxD* | 3A3L 2i *HoxD* | 0.2779 |
| WT J1 Serum Ctrl | WT J1 2i Ctrl | 0.2195 |
| WT J1 Serum Ctrl | 3B3L Serum Ctrl | 0.0513 |
| WT J1 Serum Ctrl | 3B3L 2i Ctrl | 0.1445 |
| 3B3L Serum Ctrl | 3B3L 2i Ctrl | 0.5587 |
| WT J1 Serum *Shh-Mnx1* | WT J1 2i *Shh-Mnx1* | 0.0001 |
| WT J1 Serum *Shh-Mnx1* | 3B3L Serum *Shh-Mnx1* | 0.2274 |
| WT J1 2i *Shh-Mnx1* | 3B3L 2i *Shh-Mnx1* | <0.0001 |
| 3B3L Serum *Shh-Mnx1* | 3B3L 2i *Shh-Mnx1* | 0.3214 |
| WT J1 Serum *En2-Shh* | WT J1 2i *En2-Shh* | 0.0228 |
| WT J1 Serum *En2-Shh* | 3B3L Serum *En2-Shh* | 0.4785 |
| WT J1 2i *En2-Shh* | 3B3L 2i *En2-Shh* | <0.0001 |
| 3B3L Serum *En2-Shh* | 3B3L 2i *En2-Shh* | 0.0028 |
| WT J1 Serum *En2-Mnx1* | WT J1 2i *En2-Mnx1* | 0.0101 |
| WT J1 Serum *En2-Mnx1* | 3B3L Serum *En2-Mnx1* | 0.3550 |
| WT J1 2i *En2-Mnx1* | 3B3L 2i *En2-Mnx1* | 0.0001 |
| 3B3L Serum *En2-Mnx1* | 3B3L 2i *En2-Mnx1* | 0.3190 |

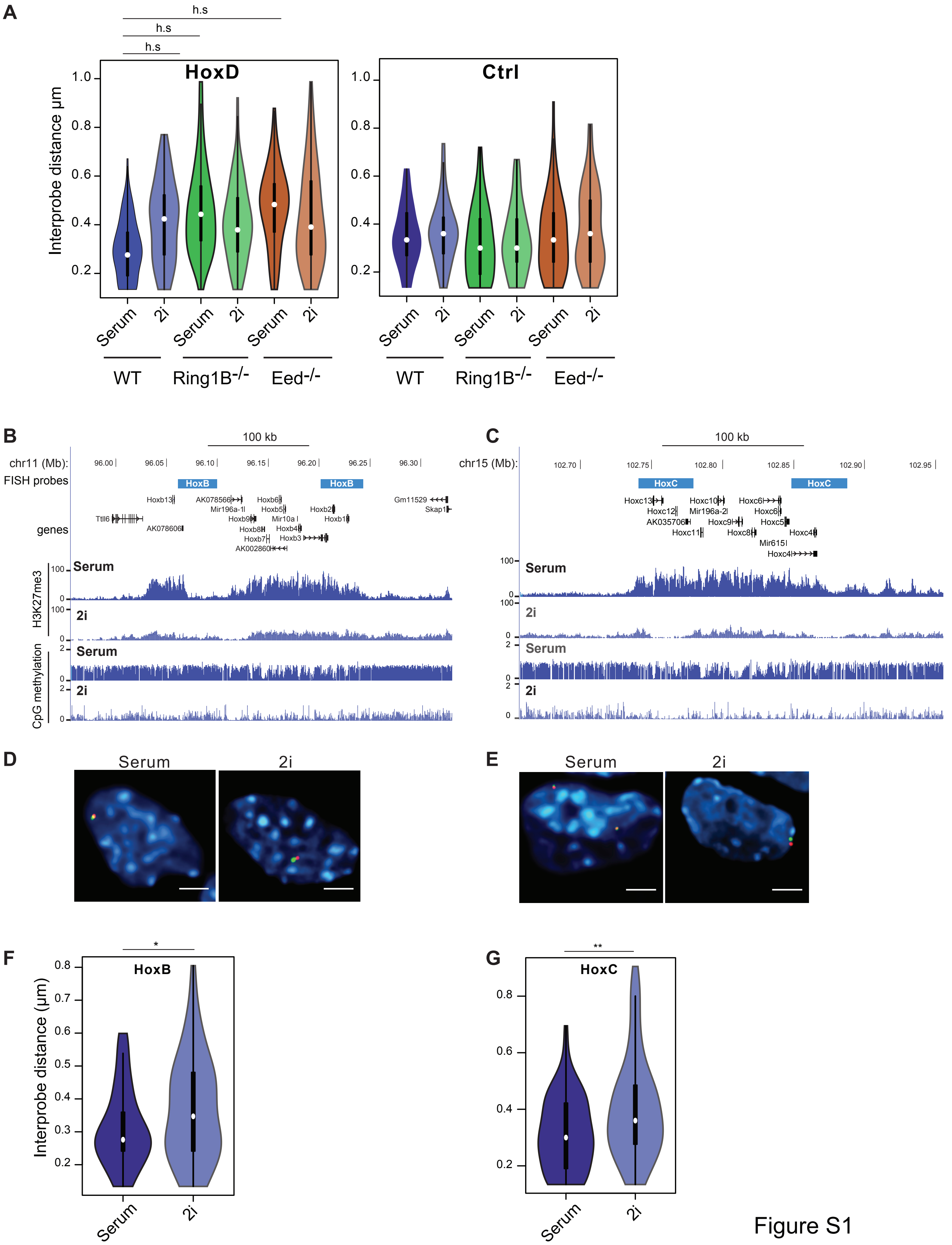
A
h.s
h.s
h.s
HoxD
Ctrl
Interprobe distance µm
1.0
0.8
0.6
0.4
0.2
Serum
2i
WT
Ring1B-/-
Eed-/-
B
100 kb
chr11 (Mb):
96.00
96.05
96.10
96.15
96.20
96.25
96.30
FISH probes
HoxB
genes
H3K27me3
CpG methylation
Serum
2i
C
chr15 (Mb):
102.70
102.75
102.80
102.85
102.90
102.95
HoxC
D
E
F
HoxB
Interprobe distance (µm)
G
HoxC
Figure S1

**Supplementary Figure 1**

A. Violin plots showing the distribution of inter-probe distances for HoxD and control (Ctrl) loci (from Figure 1A) in WT, *Ring1B*$^{-/-}$ and *Eed*$^{-/-}$ cells grown in serum or 2i. Data are abiological replicate for the data in Figure 1. h.s = p<0.0001. Details of statistical analysis are given in Tables S2 and S3.

B. UCSC genome browser tracks (mm9 assembly of the mouse genome) showing the location on chromosome 11 of FISH probes used to measure compaction across the *HoxB* locus. Probe co-ordinates are given in Supplementary Table 1. Below are shown the H3K27me3 (Marks et al., 2012) and DNA methylation (Habibi et al., 2013) profiles for this region of the mouse genome in mESCs grown in serum or 2i.

C. As in (B) but for the *HoxC* locus on chromosome 15.

D. Representative images of *HoxB* probe hybridisation signals (red and green) in WT E14 mESCs grown in serum or 2i. Scale bars represent 10 µm.

E. As in (D) but for *HoxC*.

F. Violin plots showing the distribution of inter-probe distances for the *HoxB* locus in mESCs cells grown in serum or 2i. The vertical line and spot within each plot indicate the interquartile range and median, respectively. *p<0.05, **p<0.01.

G. As in (F) but for *HoxC*.

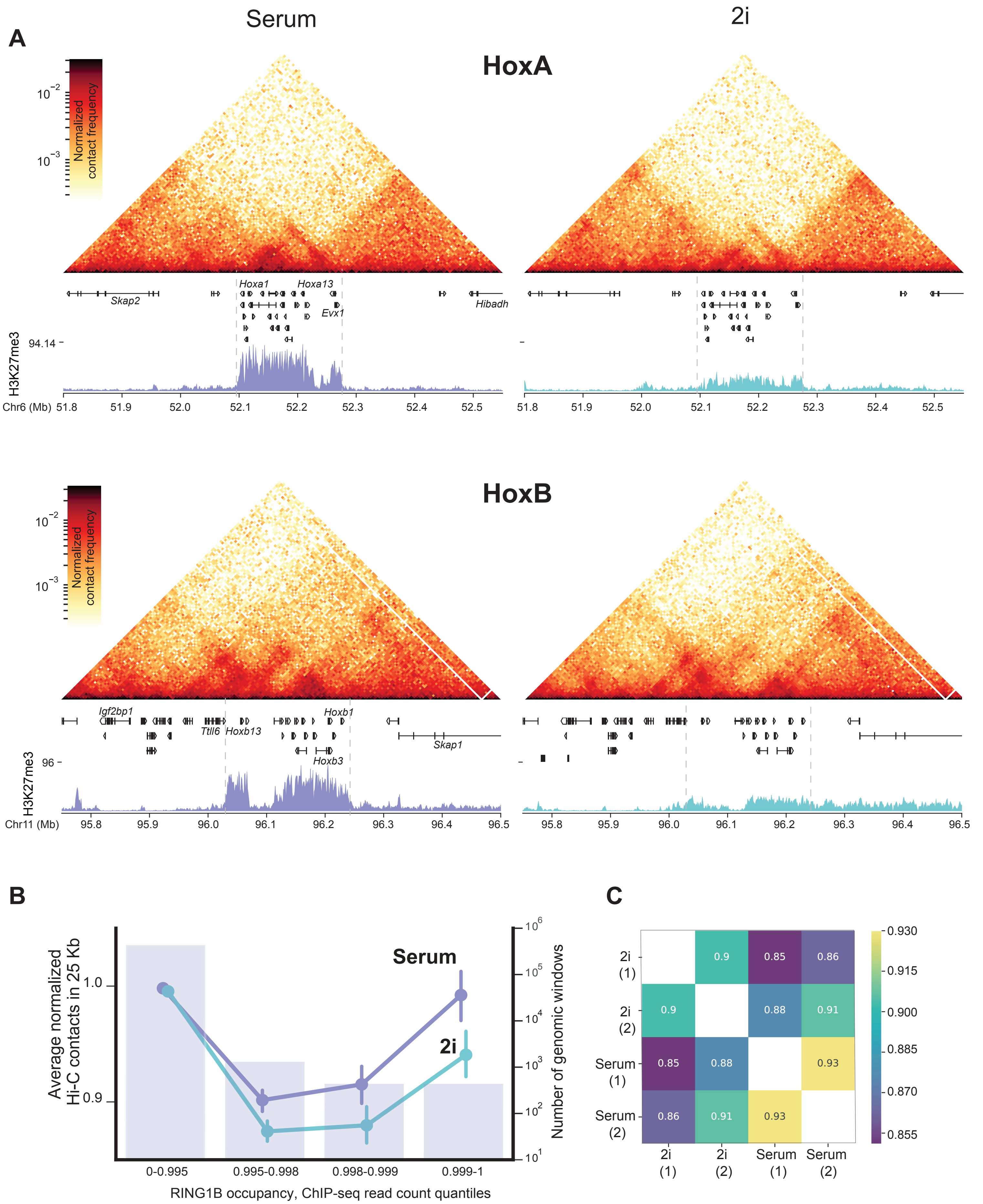
A
Serum
2i
HoxA
Normalized contact frequency
$10^{-2}$
$10^{-3}$
Hoxa1
Hoxa13
Skap2
Evx1
Hibadh
H3K27me3
94.14
Chr6 (Mb)
51.8
51.9
52.0
52.1
52.2
52.3
52.4
52.5
HoxB
Igf2bp1
Hoxb1
Ttll6
Hoxb13
Skap1
Hoxb3
96
Chr11 (Mb)
95.8
95.9
96.0
96.1
96.2
96.3
96.4
96.5
B
Average normalized Hi-C contacts in 25 Kb
Serum
2i
1.0
0.9
Number of genomic windows
$10^{6}$
$10^{5}$
$10^{4}$
$10^{3}$
$10^{2}$
$10^{1}$
0-0.995
0.995-0.998
0.998-0.999
0.999-1
RING1B occupancy, ChIP-seq read count quantiles
C
2i (1)
2i (2)
Serum (1)
Serum (2)
0.9
0.85
0.86
0.88
0.91
0.93
0.930
0.915
0.900
0.885
0.870
0.855

Figure S2

**Supplementary Figure 2**

A. Hi-C heatmaps (normalised contact frequencies) for cells grown in serum and 2i media for the *HoxA* (top) and *HoxB* (bottom) clusters along with gene annotations and H3K27me3 ChIP-seq profiles (Marks et al., 2012). Boundaries of the *Hox* clusters are marked with dashed lines.
B. Same as Fig. 3B, but with RING1B ChIP-seq quantification instead of H3K27me3 (both Hi-C data compared to RING1B data from serum-grown cells).

C. Correlation of $\log_2$ of insulation score profiles (100 kb window) across serum and 2i Hi-C replicates.

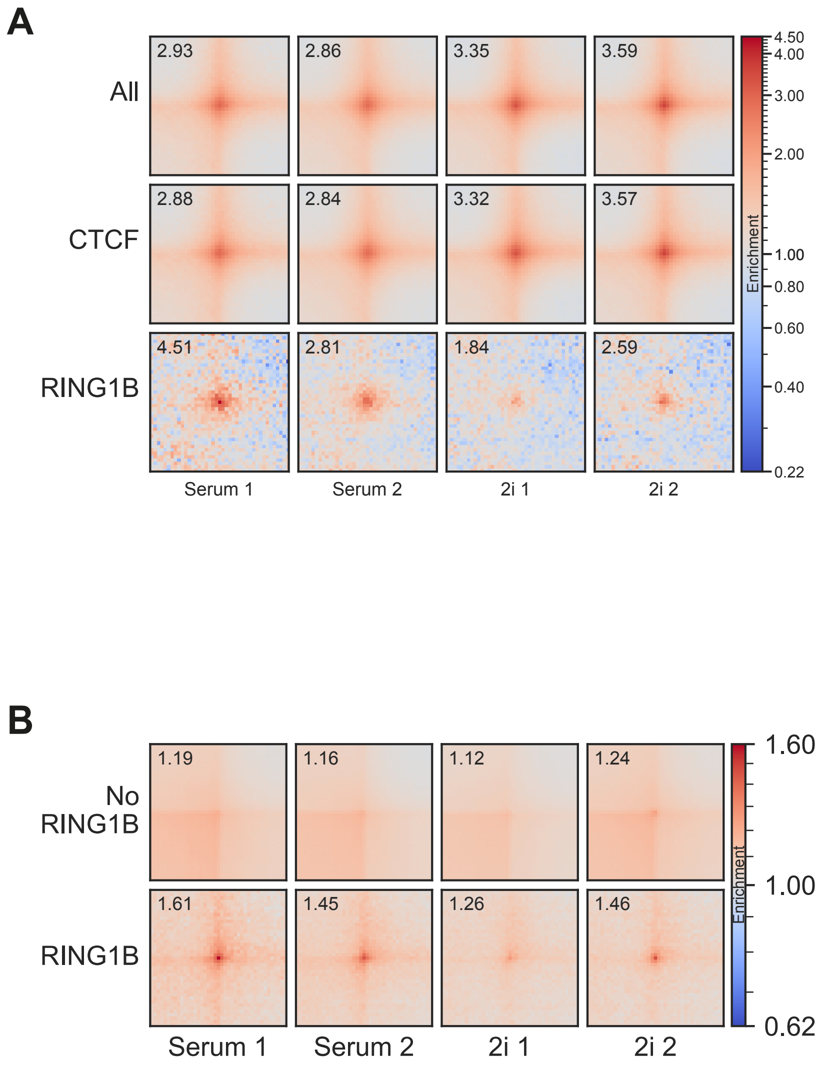

A
2.93
2.86
3.35
3.59
All
2.88
2.84
3.32
3.57
CTCF
4.51
2.81
1.84
2.59
RING1B
Serum 1
Serum 2
2i 1
2i 2
4.50
4.00
3.00
2.00
1.00
0.80
0.60
0.40
0.22
Enrichment
B
1.19
1.16
1.12
1.24
No RING1B
1.61
1.45
1.26
1.46
RING1B
Serum 1
Serum 2
2i 1
2i 2
1.60
1.00
0.62
Enrichment


Figure S3

**Supplementary Figure 3**

A. As for main Figure 4C, but for individual replicates.

B. As for main Figure 4D, but for individual replicates of our data.

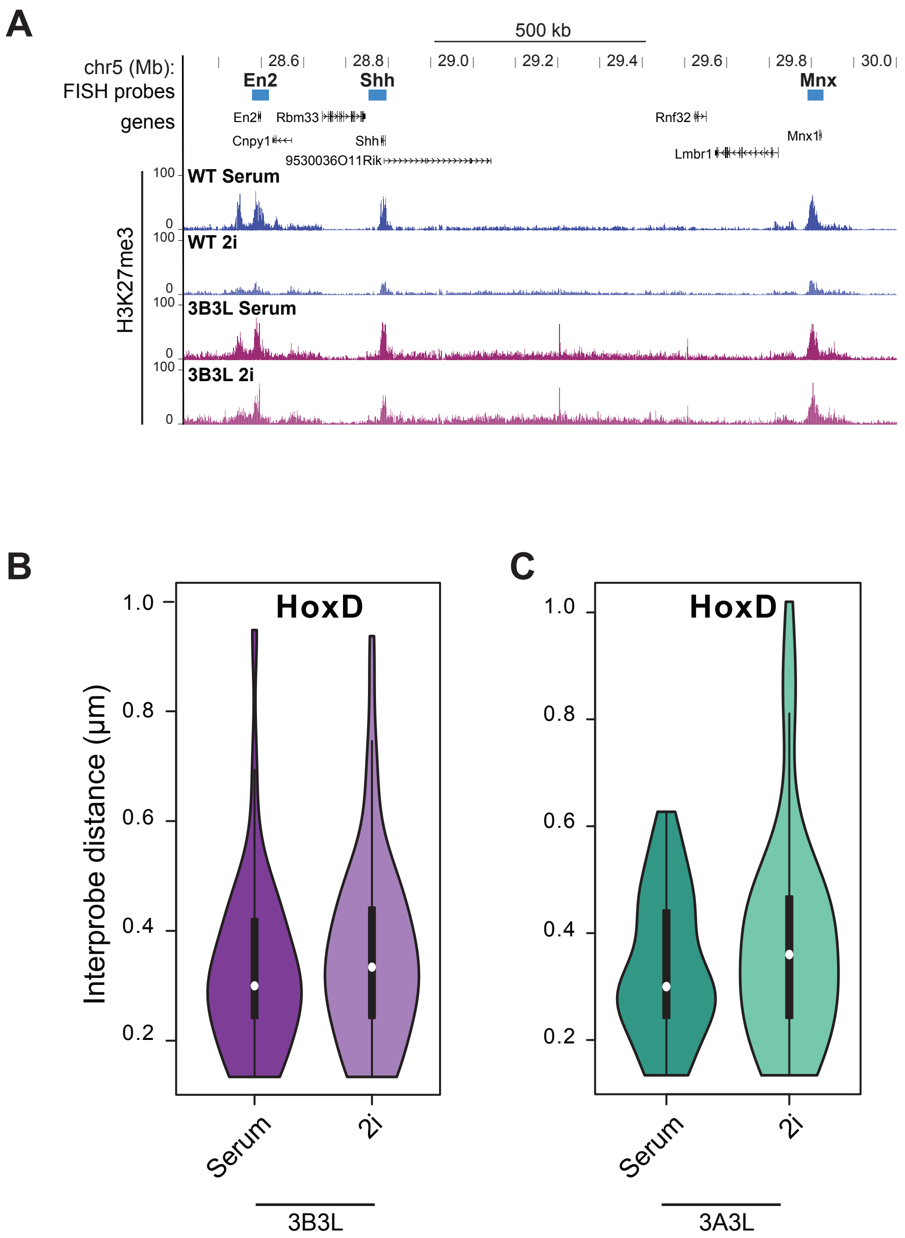
A
500 kb
chr5 (Mb):
28.6
28.8
29.0
29.2
29.4
29.6
29.8
30.0
FISH probes
En2
Shh
Mnx
genes
En2
Rbm33
Cnpy1
Shh
9530036O11Rik
Rnf32
Lmbr1
Mnx1
H3K27me3
WT Serum
WT 2i
3B3L Serum
3B3L 2i
100
0
B
HoxD
Interprobe distance (µm)
1.0
0.8
0.6
0.4
0.2
Serum
2i
3B3L
C
HoxD
Serum
2i
3A3L

Figure S4

**Supplementary Figure 4**

A. UCSC genome browser tracks (mm9 assembly) showing the location on chromosome 5 of FISH probes used to measure distal interactions across the Shh locus. Probe co-ordinates are given in Supplementary Table 1. Below are shown the H3K27me3 profiles for this region of the mouse genome in WT (Marks et al., 2012) and 3B3L mESCs grown in serum or 2i.

B. Violin plots showing distribution of inter-probe distances at the *HoxD* locus 3B3L cells cultured in serum/LIF and 2i/LIF. This is a biological replicate for the data in Figure 5E.

C. As for (A) but for 3A3L cells.
